# Directed evolution of generalist biosensors for single ring aromatics

**DOI:** 10.1101/2022.12.11.519963

**Authors:** Hannah O. Cole, Clayton W. Kosonocky, Mason Schechter, Jimmy Gollihar, Andrew D. Ellington, Simon d’Oelsnitz

**Affiliations:** Department of Molecular Biosciences, University of Texas at Austin, Austin, Texas 78712, United States; Center for Infectious Diseases, Department of Pathology and Genomic Medicine, Houston Methodist Research Institute, Houston, TX 77030, USA

**Keywords:** Biosensors, directed evolution, metabolic engineering, protein engineering

## Abstract

Biosensors can accelerate the engineering of new biosynthetic pathways. Phloroglucinol is a platform chemical of wide utility that can be produced at limited titers in *Escherichia coli*. Starting from the TetR family repressor RolR that is responsive to the related compound resorcinol, we were able to employ a combined selection and screen to identify variants that had greatly improved activities with phloroglucinol (EC50 for phloroglucinol of 131 uM, relative to an estimated 42 mM for wild-type RolR). The variants obtained were further screened with a panel of similar single ring aromatics, and several were found to be generalists, consistent with the hypothesis that both natural and directed evolution tend to first create semi-specific pockets prior to further optimization for new function

## 1. Introduction

The biomanufacture of small molecules can help to alleviate many issues associated with traditional chemical synthesis. As an example, phloroglucinol (PG), a commodity chemical, is used as a precursor for the production of a wide array of cosmetics, dyes, adhesives, explosives, and pharmaceuticals **[1-3]** yet the chemical synthesis of PG is not only costly but dangerous. Hazardous chemicals, such as the explosive trinitrotoluene (TNT), are used as the starting points for synthesis, and the downstream chemical transformations require a variety of environmentally damaging chemicals that create toxic byproducts **[4]**. The type III polyketide synthase PhlD, encoded by the phlACBDE gene cluster from *Pseudomonas putida*, can be used to condense three molecules of malonyl-CoA into PG **[5]**, but current biosynthetic titers of PG are far too low for commercial production.

While PG-producing strains can be engineered for higher titers, their development is stymied by low-throughput methods for PG measurement **[6]**. This measurement bottleneck can potentially be addressed by developing genetic biosensors that can be used for high-throughput screening, either in cells or outside of them. However, for most compounds of interest, such as PG, a corresponding biosensor is not readily available. Recent advances in transcription factor engineering have facilitated the development of novel biosensors with tailored ligand specificities **[7]**. For example, transcription factor engineering has been used to expand the binding repertoire of the transcription factor LuxR to accommodate a more diverse array of homoserine lactones **[8]**, create LacI mutants that bind new modified sugars **[9]**, and identify AraC mutants responsive to mevalonate **[10]**.

To facilitate rapid PG measurement, we have used a directed evolution method, SELIS (Seamless Enrichment of Ligand-Inducible Sensors) to generate a tractable PG biosensor. Starting from the TetR family repressor RolR, which is natively responsive to the PG analog resorcinol, we were able to alter the ligand-binding site to accommodate an additional hydroxyl and have identified several changes to the RolR binding pocket that may provide a more general route to the identification of additional biosensors for benzene derivatives.

## 2. Materials and Methods

### 2.1. Plasmid cloning & strain engineering

Plasmid cloning, directed evolution, and biosensor screens were all performed in Escherichia coli DH10B (New England BioLabs, Ipswich, MA, USA). Growth media consisted of LB-Miller (LB) media (BD, Franklin Lakes, NJ, USA) and 1.5% LB-Agar also from BD. All plasmid constructs are available in Supplementary Materials Table S1 and will be made available on Addgene. Plasmids were assembled using Gibson and Golden gate cloning methods and all DNA parts were ordered as gBlocks from IDT.

### 2.2. Benzene alcohol ligands

Screens were performed with chemicals: Resorcinol (Oakwood Chemical, CAT#: 168800), phloroglucinol (Combi-Blocks, CAT#: QC-6886), pyrogallol (Sigma Aldrich, CAT#: 254002), orcinol (Sigma Aldrich, CAT#: 44720), 1,2,4-benzenetriol (Sigma Aldrich, CAT#: 173401).

### 2.3. Chemical Transformation

All transformations were done using chemically competent cells. Competent cells were produced in house using DH10B *Escherichia coli* cells. Cells were prepared using a 5mL DH10B overnight culture which was pelleted (3500g, 4 °C, 10 min), washed in 70mL buffer (10% glycerol, 100 mM CaCl2). This wash process was then repeated, and the final pellet was resuspended in 20uL of the same buffer (10% glycerol, 100 mM CaCl2). After resting on ice for 30 minutes cells were divided into 250 μL aliquots and flash frozen in liquid nitrogen prior to being stored at −80 °C.

### 2.4. Promoter Selection

We utilized the P750 and P250 promoters characterized in previous work **[11]**. These promoter sequences were based on the J23100 Anderson constitutive promoter and were designed by taking the J23100 promoter sequence, appending an upstream activating sequence, and modifying a single base in either the -35 region (TTTACA), or the -10 region (TACAAT) thus producing a promoter series of variable expression strengths. The medium strength P250 promoter differs from J23100 by a single A → C substitution in the final base of the -35 region (TTTACA → TTTACC), and the inclusion of the sequence (AAAATATATTTTTCAAAAGTATCG) upstream from the -35 site. The strong promoter P750 differs from wild type by a T → G substitution in the -35 region (TTTACA → TTGACA), and the inclusion of the sequence (CTCAGTGGCGCGCCTCAGTCCTCG) upstream from the -35 site. For the modified Prolr promoter, the RolR operator sequence was inserted using Gibson assembly. Terminators were placed upstream of our promoter sequences to prevent interference from any upstream transcription occurring simultaneously. Additionally, we included RiboJ in each of our promoters to minimize variability between operators and increase fluorescent signal.

### 2.5. Measuring Initial RolR Biosensor Response

Our biosensors were evaluated by first chemically co-transforming our pRolR sensor production plasmid and our pORolR-GFP reporter plasmid into DH10B cells. Upon transformation and plating on LB agar with the appropriate antibiotics, colonies were picked in triplicate and cultured overnight. After overnight incubation a deep-well plate (Corning, Product #: P-DW-20-C-S) was inoculated (900 μL of LB media per well) with 20 μL of overnight co-transformation culture and sealed with an AeraSeal film (Excel Scientific, Victorville, CA, USA). Plates were left at 37 °C, 250 rpm for two hours and then induced. Induction occurred with 110 μL of LB (negative controls) or 100 μL of LB supplemented with 10 μL of resorcinol dissolved in LB. Upon induction plates were returned to incubate at 37 °C, 250 rpm for an additional 4 hours. Plates were then spun down (3500g, 4 °C, 10 min) and the supernatant removed. Cell pellets were resuspended in 1mL of PBS (137 mM NaCl, 2.7 mM KCl, 10 mM Na2HPO4, 1.8 mM KH2PO4, pH 7.4) and 100 μL of resuspended pellet was added to a 96 well plate (Corning, Product #: 3904). Fluorescence was measured on a Tecan Infinite M200 plate reader set to measure Ex: 475 nm, Em: 509 nm and absorbance 600nm.

### 2.6. RolR Library Construction

Libraries were constructed via PCR amplification of the pRolR plasmid using primers (IDT) containing three sets of NNS degenerate nucleotides targeting three separate residues. The resulting PCR reaction was DpnI digested for 16 hours, gel extracted, assembled using the Gibson method, and finally transformed into *E. coli* DH10B cells bearing the pSELIS-RolR plasmid. Transformation efficiency exceeded 10^6^, indicating several fold coverage of the library. Transformed cells were grown in LB media overnight at 37 °C in carbenicillin and chloramphenicol.

### 2.7. Directed Evolution of RolR and screening for a PG Biosensor

Our transformed RolR mutant library was grown up overnight and 20 μL of overnight was used to seed a 5mL culture containing LB and antibiotics (Chloramphenicol and Ampicillin), and 100 μg/mL zeocin (Thermo Fisher. CAT#: R25001). These cultures were grown for 7 hours at 37 °C and diluted by taking 0.5 μL of culture and adding it to 1 mL of LB. These were further diluted 100 μL of each dilution and adding it to 900 μL of LB. These final dilutions were then plated taking by 300 μL of dilution and plating on 3 different plates each containing chloramphenicol, ampicillin, and zeocin dissolved in water. After incubating overnight at 37 °C the most highly fluorescent colonies were picked and grown overnight in a deep-well plate in 1 mL and antibiotics. Plates were sealed with an AeraSeal film and were incubated at 37 °C. We inoculated 5 mL of LB with a glycerol stock of our pSELIS-RolR and pRolR and grew this overnight.

The following day, 20 μL of each culture was used to inoculate two separate wells within a new 96-deep-well plate containing 900 μL of LB media. Additionally, eight separate wells containing 1 mL of LB media were inoculated with 20 μL of the overnight culture expressing the parental RolR variant. A typical arrangement would have 44 unique clones on the top half of the plate, duplicates of those clones on the bottom half of the plate, and the rightmost column occupied by cells harboring the parental RolR variant. After 2 h of growth at 37 °C the top half of the 96-well plate was induced with 100 μL of LB media whereas the bottom half of the plate was induced with 100 μL of LB media containing the target phloroglucinol dissolved in 10 μL of water. The concentration of phloroglucinol used for induction is typically the same concentration used in the LB agar plate for screening during that particular round of evolution. Cultures were grown for an additional 4 h at 37 °C, 250 rpm and subsequently centrifuged (3500g, 4 °C, 10 min). Supernatant was removed and cell pellets were resuspended in 1 mL of PBS. One hundred μL of the cell resuspension for each condition was transferred to a 96-well microtiter plate, from which the fluorescence (Ex: 475 nm, Em: 509 nm) and absorbance (600 nm) was measured using the Tecan Infinite M200. Clones with the highest signal-to-noise ratio were then sequenced and subcloned into a fresh pRolR vector.

For sensor variant dose response measurement, each RolR variant plasmid was co-transformed into DH10B cells alongside pORolR-GFP and three individual colonies from each transformation were subsequently grown overnight. The resulting cultures were then assayed, as described in Biosensor Response Measurement, using 11 different concentrations of the target phloroglucinol alongside an LB-only control.

### 2.8. AI-Assisted Molecular Docking

Predicted structures for the wild-type and mutant RolR structures were generated with the AlphaFold Multimer Google Collaboratory notebook **[12]**. A comparison of the wild-type RolR structure and the AlphaFold-predicted RolR structure is shown in **Supplementary Figure 1**. These structures were then passed into GNINA with the ligand structures of resorcinol and phloroglucinol to obtain predicted dockings positions **[13]**. The GNINA parameters used were exhaustiveness of 64, no convolutional neural network scoring, random seed, and auto-box ligand set to the entire protein. Each time GNINA was run, 20 different docking positions were outputted, ranked by energetics of the ligand-protein fit. Some of the predicted docking positions were not plausible, so predicted docking configurations in the vicinity of the known resorcinol-binding pocket were chosen. Docked structures were then visualized and analyzed using UCSF ChimeraX **[14-15]**.

## 3. Results

### 3.1. Domestication of a resorcinol catabolism regulator

To identify a biosensor that could recognize phloroglucinol, we sought regulators responsive to ligands that were chemically similar to our target compound, in particular benzenetriols. The transcription factor RolR controls resorcinol catabolism in *Corynebacterium glutamicum*, has previously been found to bind resorcinol which differs from phloroglucinol by a single hydroxyl moiety (Figure 1a), and thus was potentially an excellent starting point for directed evolution.

**Figure 1.**
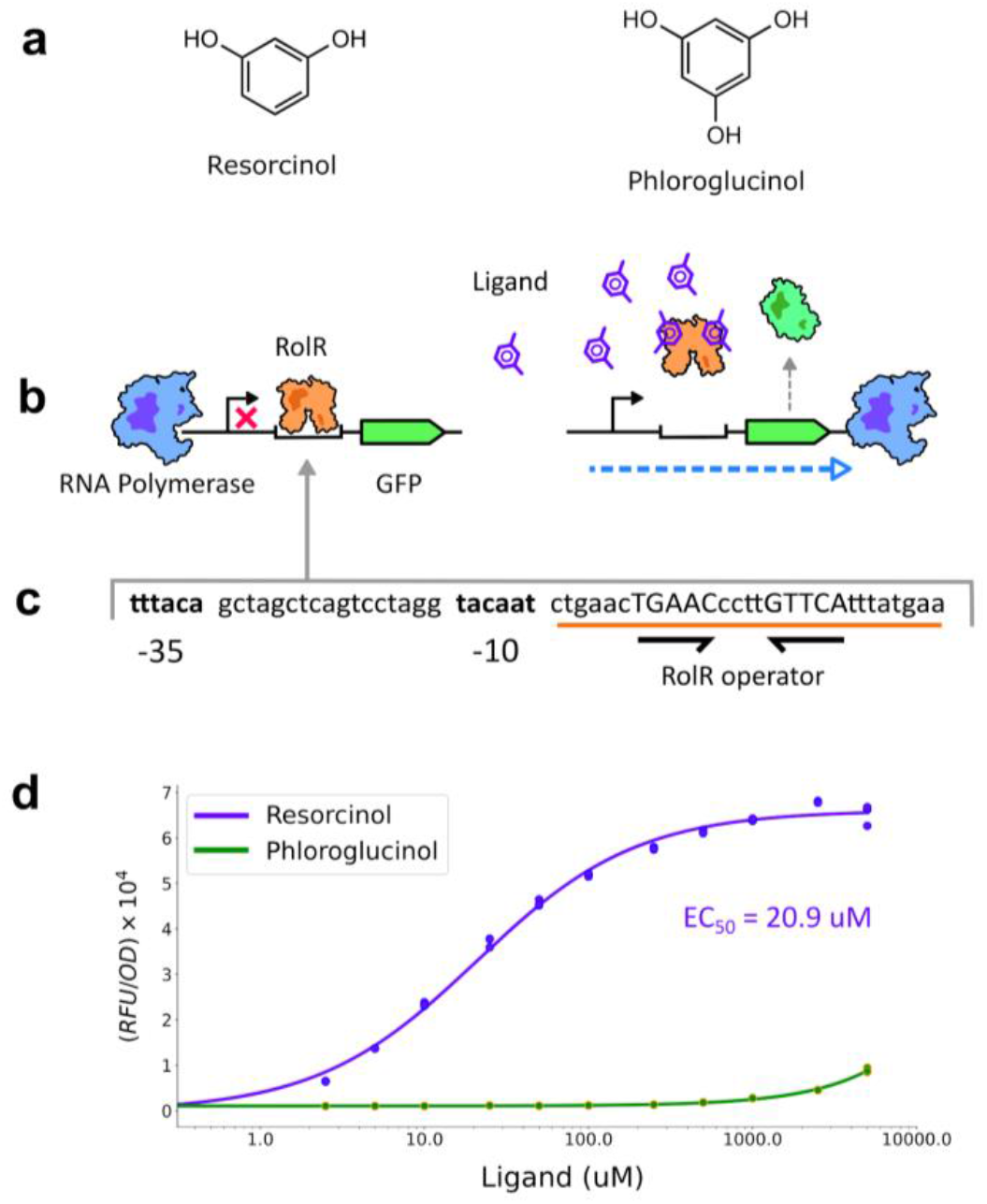
Domestication of the RolR transcription regulator as a resorcinol-responsive biosensor. **(A)** Resorcinol and phloroglucinol chemical structures. **(B)** Schematic of the synthetic RolR-regulated genetic circuit. **(C)** Sequence of the pORolR promoter. **(D)** Response of the RolR-regulated circuit to resorcinol and phloroglucinol. Measurements represent the average of three biological replicates.

A RolR expression construct (pRolR) was generated in which the protein was expressed by a variant of the constitutive Anderson series promoters J23100 (P250) in which the sequence of the -35 region was modified with an A → C base change (TTTACA → TTTACC), a mutation that had previously been shown to reduce expression strength **[11]**. A reporter plasmid (pORolR-GFP) contained the sfGFP gene under the control of a stronger synthetic promoter, P750, upstream from the Orolr operator sequence identified from the literature **[16]** (**Figure 1b** and **1c**). The choice of the relative strengths of the promoters was intended to allow the observation of a maximal dynamic range for RolR-based induction. The expression and reporter constructs were assayed together in the presence of varying amounts of resorcinol and phloroglucinol. Initial assays showed a concentration-dependent fluorescent signal from 2.5 uM to 5000 uM of its native ligand resorcinol, with an EC50 of 21 uM and a 60-fold increase in fluorescent signal observed at the highest concentrations tested (5000 uM) (**Figure 1d**). Initial responsivity to phloroglucinol indicated that directed evolution could likely be used to improve upon nascent interactions and activation.

### 3.2. Evolution of a phloroglucinol responsive biosensor

We have previously developed a conjoined selection and screening method that we call SELIS (Seamless Enrichment of Ligand Inducible Sensors) **[11]**. In this method an inverter circuit activates the expression of an antibiotic resistance element in the absence of an inducer, thus allowing for the selection of active repressor variants that interact with their operator. Following the down-selection of only active repressor variants, survivors are plated in the presence of an inducer, and those colonies that show activation of GFP expression are picked for further validation of ligand response.

Mutant RolR libraries were then generated via PCR using primers containing NNS degenerate codons targeted to residues 110, 111, and 114 (Figure 2) because they were proximal to the carbon at which phloroglucinol carries an additional hydroxyl group and had been shown to be directly involved in resorcinol ligand binding **[16]**. The mutant library spanning ca. 8,000 (20×20×20) protein variants was selected on zeocin, and subsequently screened for GFP induction in the presence of 4 mM phloroglucinol. The library was quickly winnowed to only three mutants (A9, B7, and C7) that showed significant improvement in phloroglucinol binding over wild-type RolR. Mutant A9 was found to contain a single mutation (L114T), while B7 and C7 contain two mutations each: (L111V, L114V) and (L111C, L114M), respectively.

**Figure 2.**
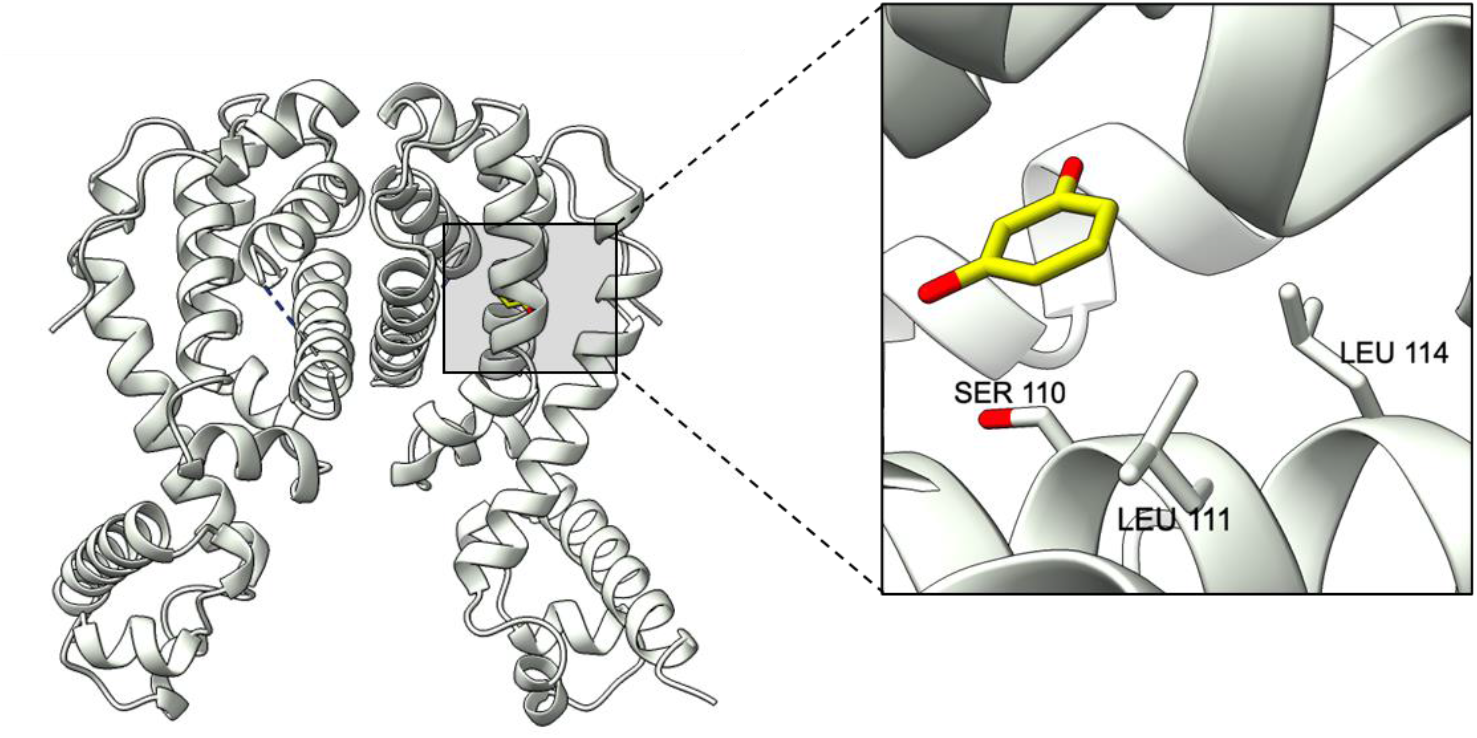
RolR library design. The structure of RolR solved in complex with resorcinol (PDB: 3AQT) was used to guide library design. The residues S110, L111, and L114 were selected for site saturation mutagenesis based on their proximity to the carbon of resorcinol where phloroglucinol bears a hydroxyl group.

### 3.3. Evolution of generalist sensors

Each of the variants were assayed for response to the original ligand, resorcinol, and to the challenge ligand phloroglucinol (Figure 3a). All variants retained activity with resorcinol, with A9 and B7 showing an improved response (Figure 3b). In addition, all three variants showed improvements with the target effector, phloroglucinol. In particular, A9 showed a dramatically improved response, with an EC50 for phloroglucinol of 131 uM (relative to an estimated 42 mM for wild-type RolR). Thus, the A9 variant in particular appears to be an effector generalist, capable of responding to similar concentrations of both resorcinol (17uM) and phloroglucinol (131 uM).

**Figure 3.**
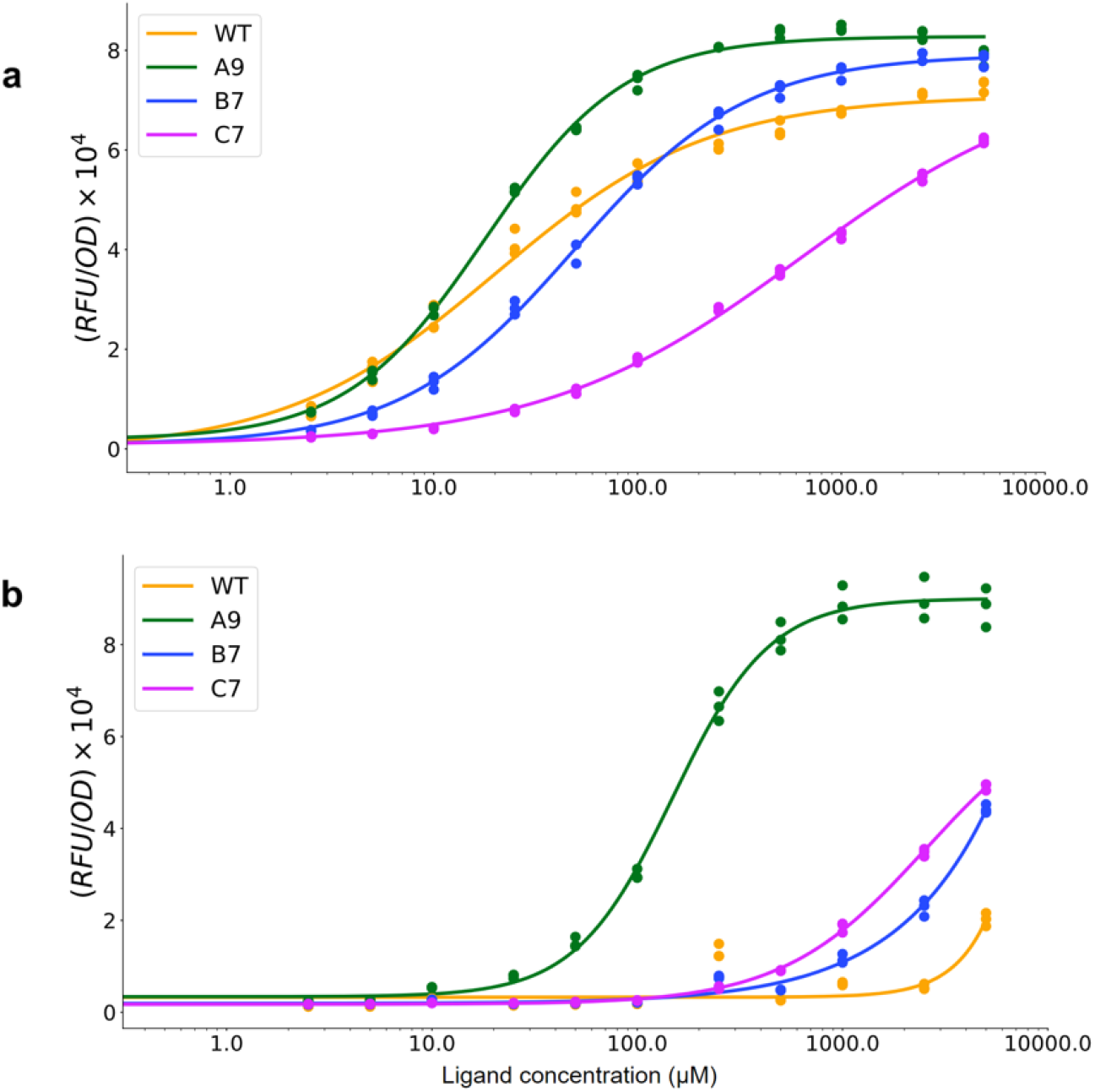
RolR variant response to resorcinol and phloroglucinol. **(A)** RolR variant response to resorcinol. **(B)** RolR variant response to phloroglucinol. Measurements for each condition represent the average of three biological replicates.

### 3.4. Rationalizing and predicting effector specificity

The ability to model small ligand binding to proteins has undergone revolutionary improvements **[17-18]**, especially with the development of sophisticated software packages such as AlphaFold for structure determination **[19]** and GNINA for docking **[20]**. We therefore attempted to rationalize the results obtained so far by modeling the different variants using AlphaFold to predict the structures of both wild-type and mutant proteins, and GNINA to dock phloroglucinol into the known effector-binding site. As has been seen in many other studies, the AlphaFold structure matched up almost exactly with the known crystal structure of RolR (**Supplementary Figure 1**). GNINA docking of phloroglucinol into the effector binding site of the wild-type structure yielded the same pose as GNINA-docked resorcinol (**Supplementary Figure 2**). However, when phloroglucinol was docked into the AlphaFold-predicted structures, the A9 variant adopted a shifted ligand pose compared to the wild-type or the other variants (**Figure 4**), with the L114T substitution potentially donating a new hydrogen bond to the ligand (**Figure 4b**).

**Figure 4.**
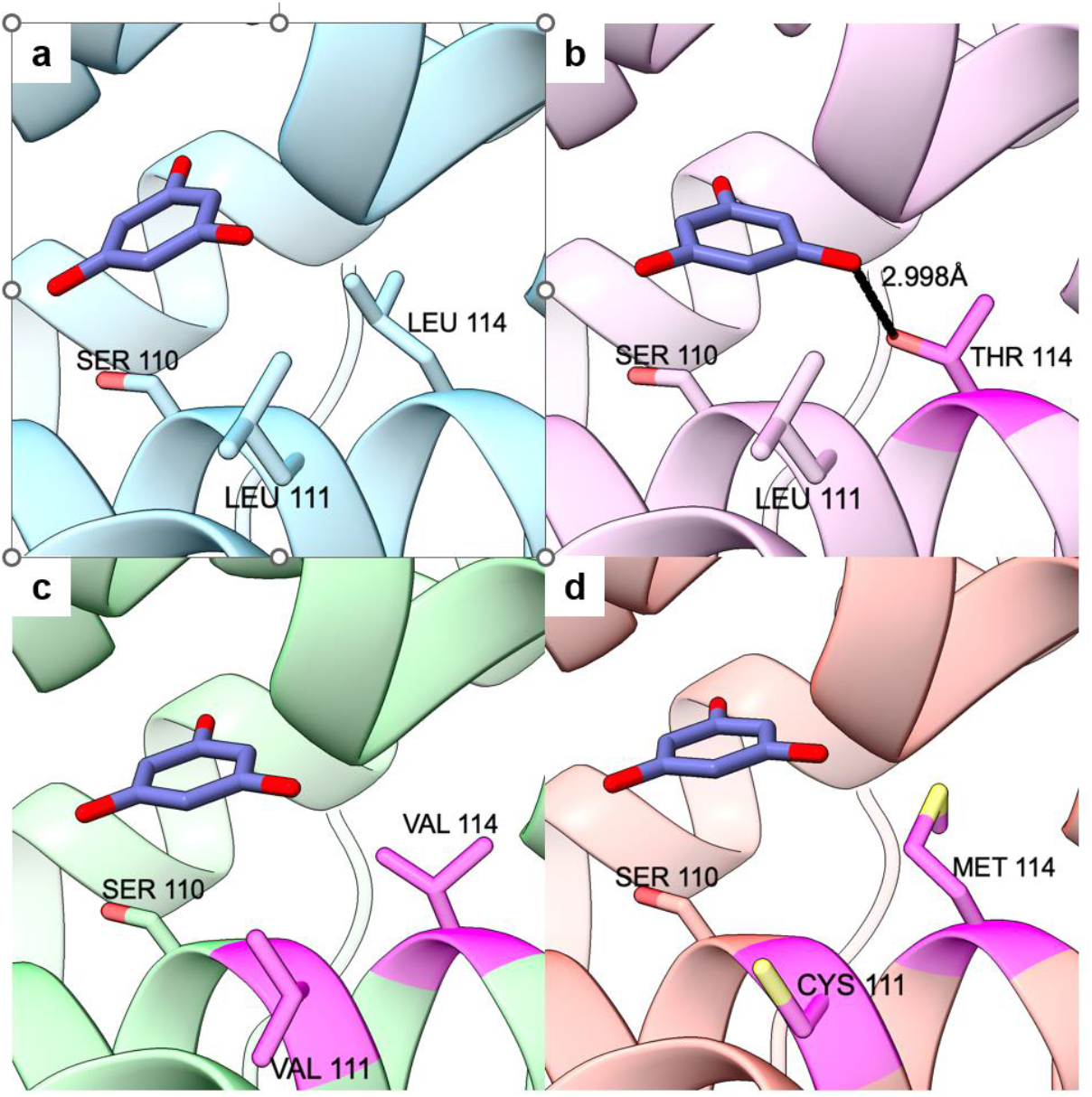
Comparison of GNINA-docked phloroglucinol positions on AlphaFold-predicted RolR wild-type and mutant structures. Mutated residues shown in dark pink. **(A)** WT RolR, **(B)** Mutant A9 (L114T), **(C)** Mutant B7 (L111V, L114V), **(D)** Mutant C7 (L111C, L114M)

### 3.5. The bounds of effector specificity

Based in part on the computational analyses carried out, we hypothesized that the RolR generalist provided a basis for moving effector specificity in a number of directions, beyond binding to phloroglucinol. To test this hypothesis, we screened wild-type RolR and its variants (A9, B7, and C7) with a panel of benzene triol compounds that were structurally related to both resorcinol and phloroglucinol (**Figure 5**). Wild-type RolR displays high specificity for resorcinol ((11); **Figure 5b**). The other variants are responsive to resorcinol, phloroglucinol, and orcinol at various levels. The expansion and rearrangement of the pocket to accommodate phloroglucinol (**Figure 4**) also allows accommodation of the methyl group of orcinol; in particular, the retraction of leucine 114 to threonine (A9) or valine (B7) provides room for the ortho methyl group. Interestingly, B7 favors orcinol over phloroglucinol, while for A9 this situation is reversed. This may be a result of the fact that the hydrophobic interactions between the two valine substitutions and orcinol are favored in B7, relative to the new hydrogen bond with phloroglucinol formed in A9. Overall, the opened pocket accommodated ligands that had functional groups that were ortho to one another, similar to the directed evolution target phloroglucinol, but not ligands that had functional groups that were meta to or para to one another (pyrogallol; 1,2,4 benzene triol).

**Figure 5.**
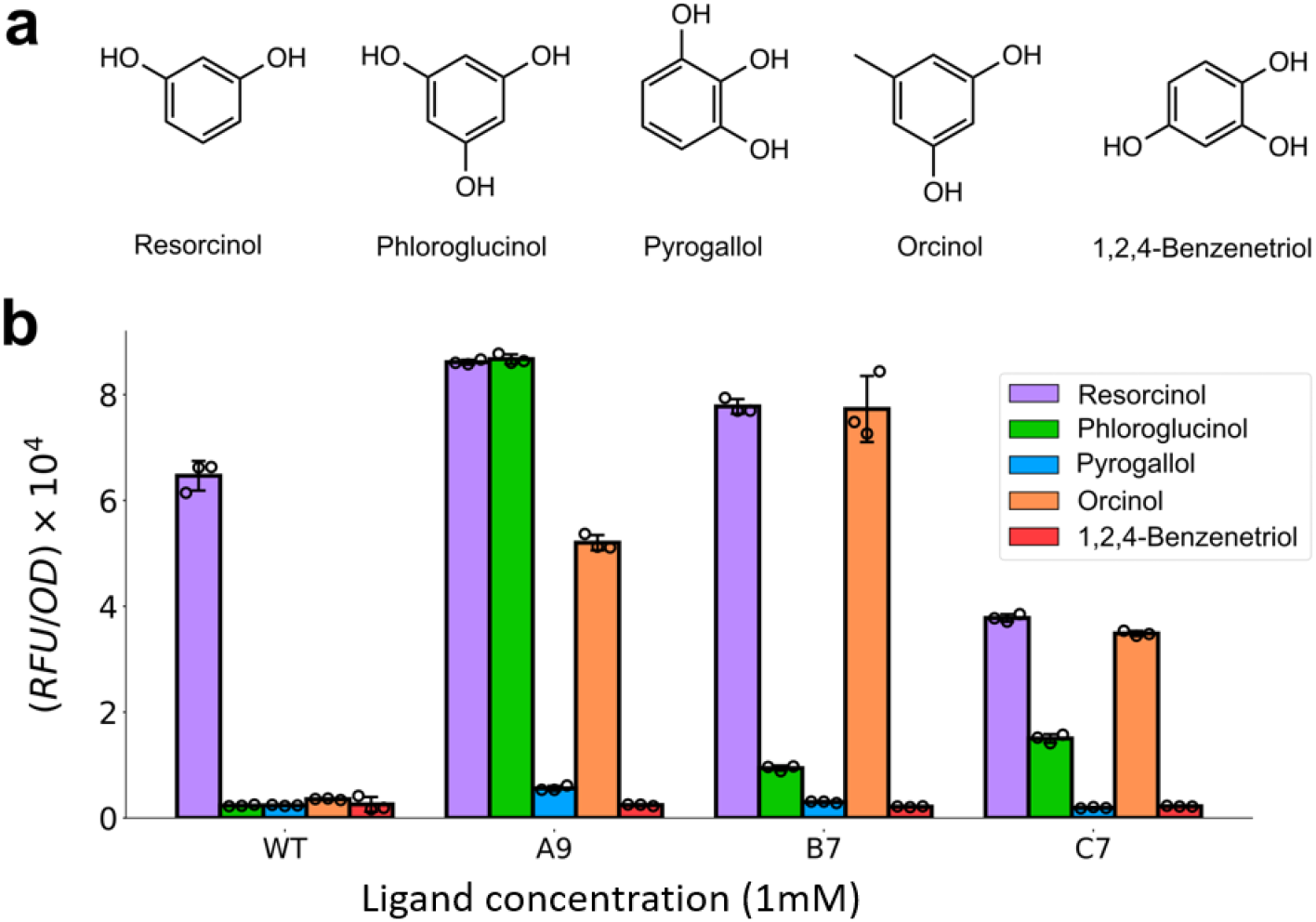
RolR Mutant Specificity. **(A)** Chemical structures used in RolR mutant screening. **(B)** RolR variant response with the native ligand resorcinol, the challenge ligand phloroglucinol, and off target benzene alcohols. Individual fluorescent measurement values for each condition performed in biological triplicate are displayed. Error bars represent mean ± s.e.m.

## 4. Discussion

While we have previously demonstrated that a combined selection and screen, SELIS, can readily identify transcription factors with altered effector specificities, the degree to which effector specificity can be optimized by simply altering the binding pocket was unclear: in one case, we were able to use several rounds of SELIS to identify highly specific biosensors for benzylisoquinoline alkaloids, starting from RamR **[21]** ; in another case, we were able to identify generalist sensors for monoterpenes, starting from a transcription factor responsive to camphor, CamR, but did not identify CamR variants that had orthogonal specificities. The current studies used a different starting point, RolR, which is known to be responsive to resorcinol, challenged it with a structurally related ligand, phloroglucinol, and then used SELIS to assess the specificities for untargeted, ligand analogs.

These studies can also be viewed from the larger context of how evolution, whether natural or directed, can traverse fitness landscapes to identify new substrate specificities. For decades we have understood that when proteins evolve towards new substrates, they often end up revisiting evolutionary ancestors and retracing evolutionary paths in the process **[22]**. Numerous studies have suggested that in general evolution proceeds along a path from specialist to generalist and eventually to a new specialist **[23-26]**. Similarly, directed evolution experiments have shown that the first step in becoming a new specialist is often to generalize, although not at the expense of binding to the original ligand **[27-28]**.

In the progression towards the acquisition of a new specificity, a single amino acid substitution is often sufficient to greatly change enzyme stereospecificity or protein-ligand binding **[29]**. For example, attempts to engineer the already highly promiscuous γ3-humulene synthase to synthesize novel sesquiterterpenes established that a single amino acid substitution at one of the core plasticity residues (W315P) led to a more promiscuous synthase with less regioselectivity. Synthase libraries targeting the W315 residue and other residues were further screened to identify additional substitutions that led to new specificities and identified a triple mutant that could shift regioselectivity by up to a 1000-fold.

In the current work, the A9 variant of RolR contained only a single amino acid substitution, and gained the ability to bind a new effector, phloroglucinol. However, the variant was a generalist that could bind both compounds, similar to our previous directed evolution studies with (CamR), where a single substitution (A139T) led to better binding to four different monoterpenes **[11]**. When off-target ligands were examined, A9 and other variants were able to bind several other, untargeted, benzene derivatives, such as orcinol. By simply opening the binding pocket in RolR we were able to change a highly specific repressor into an ortho-benzene generalist.

Overall, the varying results that have been obtained with SELIS and other directed evolution methods can be reconciled with the large body of knowledge obtained via natural selection in two ways: first, we recapitulate the notion that the first steps towards acquiring new substrate specificities occur via gross morphological changes in the binding pocket that lead to generalist phenotypes. Second, it is likely that the larger a ligand is, the more opportunities there will be to ‘dial in’ specificity via multiple amino acid changes, which is why biosensors for specific benzylisoquinoline alkaloids were more readily acquired than biosensors for monoterpenes or benzene derivatives.

From a biotechnological perspective, as we gain understanding of metabolic pathways and explore the biosynthesis of new compounds the use of biosensors will continue to be of increasing importance. Facile engineering of new sensors will be key in our ability to meet these challenges and build robust production strains for biosynthesis. This work reaffirms that highly sensitive biosensors can be generated with minimal perturbation of the wild-type protein sequence (and likely structure). Construction and testing of additional mutant libraries and screening of those libraries with a wider variety of benzene derivatives should enable us to better understand and eventually predict amino acid substitutions for even more diverse ligands.

## Supplementary Materials

T, Figure S1: RolR WT vs. AlphaFold Predicted ; Figure S2: RolR WT vs. GNINA Predicted; Table S1: Plasmid maps for RolR two plasmid sensor.

## Author Contributions

S.D. designed experiments, and H.O.C. and M.S. performed the experiments. C.W.K. performed computational structure prediction and docking analysis. The manuscript was written by H.O.C. with support from S.D, J.G, and A.D.E. S.D and A.D.E. supervised all aspects of the study.

## Funding

This work was supported in part by a Cooperative Agreement between the University of Texas at Austin and DEVCOM Army Research (W911NF-17-2-0091) and grants from the National Institute of Standards and Technology (70NANB21H100), Air Force Office of Scientific Research (FA9550-14-1-0089), National Institutes of Health (5R01EB026533-04), and the Welch Foundation (F-1654).

## Data Availability Statement

Relevant plasmids were submitted to Addgene. Code used to generate figures for data visualization is available at https://github.com/simonsnitz/plotting.

## Conflicts of Interest

The authors declare the following competing financial interest(s): S.D. is the founder of Retna Bio, a company that commercializes genetic biosensors.

## Notes

https://github.com/simonsnitz/plotting

