## Supplemental Files 1 for "Directed evolution of generalist biosensors for single ring aromatics"

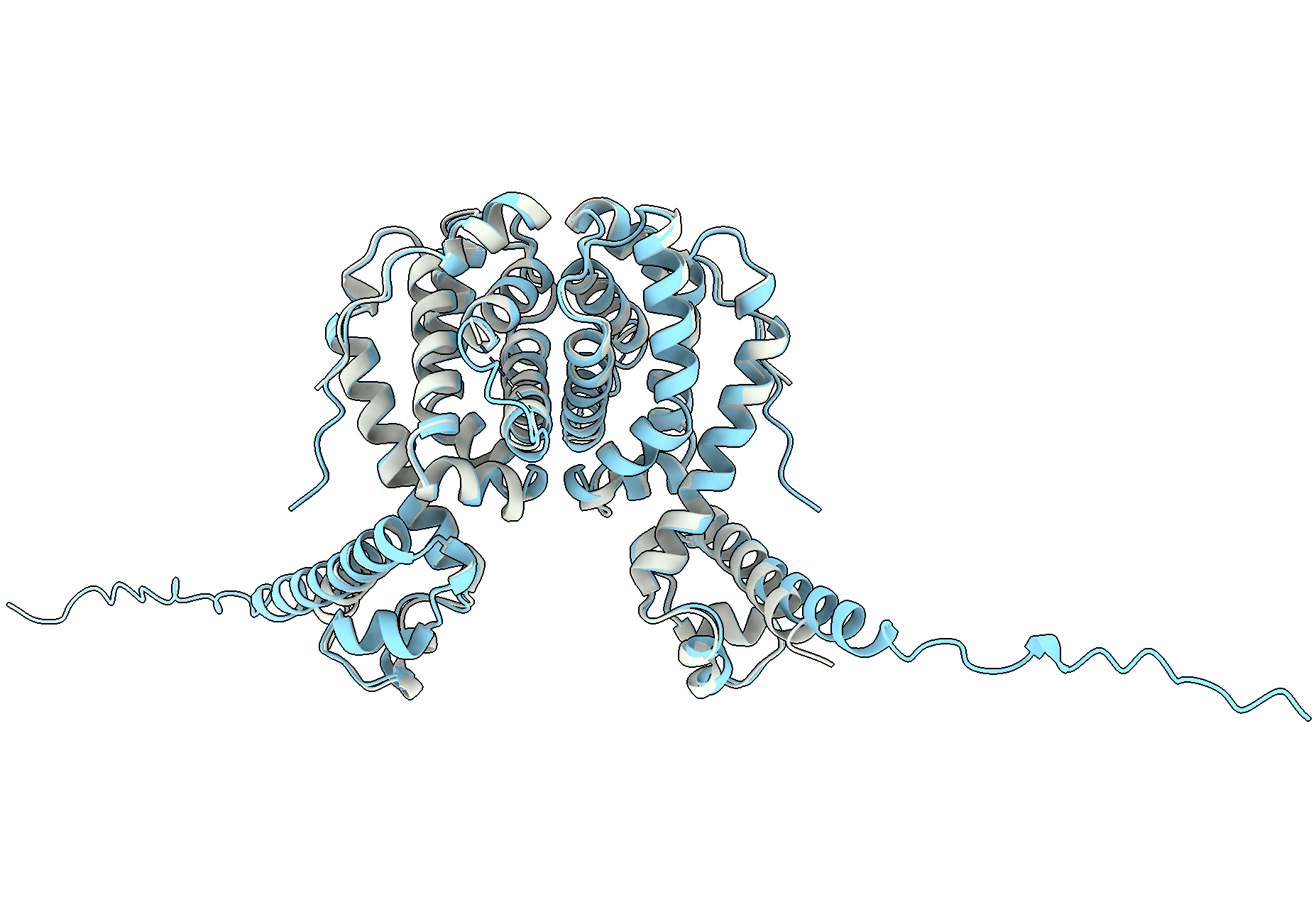

**Supplemental Figure 1.** RolR WT vs. AlphaFold Predicted. Comparison of AlphaFold-predicted RolR structure (sky blue) vs actual RolR structure (white).

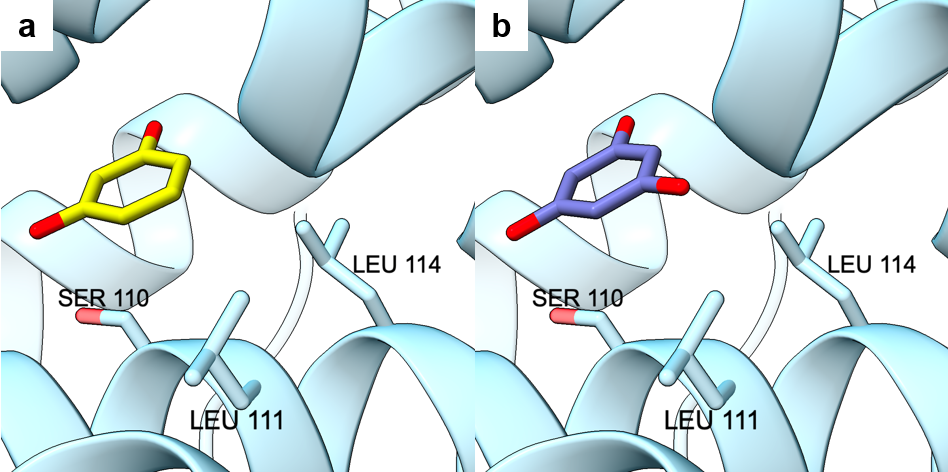

**Supplemental Figure 2.** Comparison of GNINA-docked ligand positions on AlphaFold-predicted. **(A)** Resorcinol, **(B)** Phloroglucinol

Supplementary Table 1

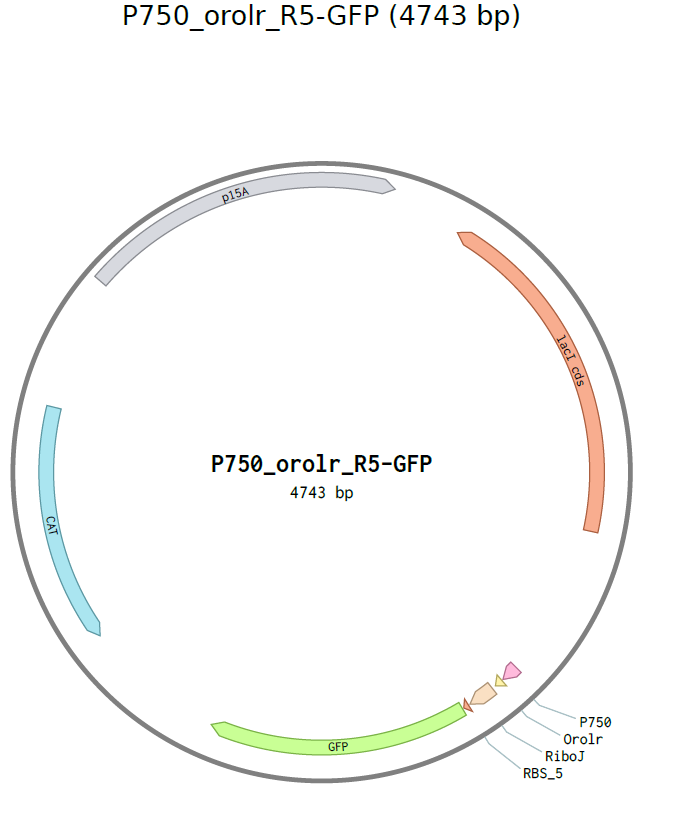

GGTTAAGGCTAAACTGAAAGGACAAGTTTTGGTGACTGCGCTCCTCCAAGCCAGTTACCTCGGTTCAAAGAGTTGGTAGCTCAGAGAACCTTCGAAAAACCGCCCTGCAAGGCGGTTTTTTCGTTTTCAGAGCAAGAGATTACGCGCAGACCAAAACGATCTCAAGAAGATCATCTTATTAATCAGATAAAATATTTCTAGATTTCAGTGCAATTTATCTCTTCAAATGTAGCACCTGAAGTCAGCCCCATACGATATAAGTTGTAATTCTCATGTTAGTCATGCCCCGCGCCCACCGGAAGGAGCTGACTGGGTTGAAGGCTCTCAAGGGCATCGGTCGAGATCCCGGTGCCTAATGAGTGAGCTAACTTACATTAATTGCGTTGCGCTCACTGCCCGCTTTCCAGTCGGGAAACCTGTCGTGCCAGCTGCATTAATGAATCGGCCAACGCGCGGGGAGAGGCGGTTTGCGTATTGGGCGCCAGGGTGGTTTTTCTTTTCACCAGTGAGACGGGCAACAGCTGATTGCCCTTCACCGCCTGGCCCTGAGAGAGTTGCAGCAAGCGGTCCACGCTGGTTTGCCCCAGCAGGCGAAAATCCTGTTTGATGGTGGTTAACGGCGGGATATAACATGAGCTATCTTCGGTATCGTCGTATCCCACTACCGAGATATCCGCACCAACGCGCAGCCCGGACTCGGTAATAGCGCGCATTGCGCCCAGCGCCATCTGATCGTTGGCAACCAGCATCGCAGTGGGAACGATGCCCTCATTCAGCATTTGCATGGTTTGTTGAAAACCGGACATGGCACTCCAGTCGCCTTCCCGTTCCGCTATCGGCTGAATTTGATTGCGAGTGAGATATTTATGCCAGCCAGCCAGACGCAGACGCGCCGAGACAGAACTTAATGGGCCCGCTAACAGCGCGATTTGCTGGTGACCCAATGCGACCAGATGCTCCACGCCCAGTCGCGTACCATCTTCATGGGAGAAAATAATACTGTTGATGGGTGTCTGGTCAGAGACATCAAGAAATAACGCCGGAACATTAGTGCAGGCAGCTTCCACAGCAATGGCATCCTGGTCATCCAGCGGATAGTTAATGATCAGCCCACTGACGCGTTGCGCGAGAAGATTGTGCACCGCCGCTTTACAGGCTTCGACGCCGCTTCGTTCTACCATCGACACCACCACGCTGGCACCCAGTTGATCGGCGCGAGATTTAATCGCCGCGACAATTTGCGACGGCGCGTGCAGGGCCAGACTGGAGGTGGCAACGCCAATCAGCAACGACTGTTTGCCCGCCAGTTGTTGTGCCACGCGGTTGGGAATGTAATTCAGCTCCGCCATCGCCGCTTCCACTTTTTCCCGCGTTTTCGCAGAAACGTGGCTGGCCTGGTTCACCACGCGGGAAACGGTCTGATAAGAGACACCGGCATACTCTGCGACATCGTATAACGTTACTGGTTTCACATTCACCACCCTGAATTGACTCTCTTCCGGGCGCTATCATGCCATACCGCGAAAGGTTTTGCGCCATTCGATGGTGTCCGGGATCTCGACGCTCTCCCTTATGCGACGCGGCCGCGAAGACACTGTCCTCAATGGTTCGTTGTGATGGCGGTAGGAATGTAATCGTTAATCCGCAAATAACGTAAAAACCCGCTTCGGCGGGTTTTTTTATGGGGGGAGTTTAGGGAAAGAGCATTTGTCATCCCGTTGAATATGGCTCGCATCTTATCGAGCATACTATCACGTCGGCGACCACTAGTCAGTTAACGCTCAGTGGCGCGCCTCAGTCCTCGTTGACAGCTAGCTCAGTCCTAGGTACAATCTGAACCCTTGTTCATTTATGAATGGCTGAGCACAGCTGTCACCGGATGTGCTTTCCGGTCTGATGAGTCCGTGAGGACGAAACAGCCTCTACAAATAATTTTGTTTAAACTAGTGAACCACGAGGCCTACATATGTCGAAGGGAGAGGAATTGTTTACTGGAGTCGTCCCAATTCTTGTTGAGTTGGACGGAGATGTCAATGGTCATAAGTTTAGCGTATCCGGTGAGGGTGAGGGAGATGCTACTTACGGAAAATTAACATTAAAATTCATTTGTACCACAGGCAAACTTCCTGTTCCTTGGCCAACTCTTGTGACGACTTTTGGCTACGGTGTGCAGTGTTTCGCTCGTTATCCTGACCACATGAAGCAACATGATTTCTTCAAGTCTGCGATGCCTGAAGGATATGTTCAAGAACGTACCATCTTCTTCAAAGATGATGGTAACTATAAGACTCGTGCAGAGGTAAAATTCGAGGGAGATACGTTGGTAAATCGCATTGAGCTGAAGGGAATCGATTTCAAGGAAGATGGAAACATTCTGGGACACAAGCTGGAGTACAATTACAATAGCCATAACGTCTATATCATGGCAGATAAGCAAAAAAACGGAATCAAGGTTAACTTCAAGATTCGCCATAATATTGAGGACGGCTCTGTGCAATTGGCGGATCATTATCAACAGAACACCCCGATTGGAGATGGTCCCGTGCTGCTGCCAGATAATCACTATTTGTCTACACAATCGGCCCTTTCCAAAGATCCGAACGAAAAGCGCGATCATATGGTACTTTTAGAGTTCGTAACTGCCGCTGGAATCACCCACGGCATGGATGAGTTGTATAAGTAATAATCCAAACCTGTTATATGTTAGCTGAGACTAGTTGGAAGTGTGGCTGTCCTCAAGCGTTTTAGTTCGTCGGTCAGTTTCACCTGATTTACGTAAAAACCCGCTTCGGCGGGTTTTTGCTTTTGGAGGGGCAGAAAGATGAATGACTGTCGGCCATTCGATGGTGTCGGGTAGCATAACCCCTTGTGATAGTCTTCGCGGCCGCTCACACTGCTTCCGGTAGTCAATAAACCGGTAAACCAGCAATAGACATAAGCGGCTATTTAACGACCCTGCCCTGAACCGACGACCGGGTCGAATTTGCTTTCGAATTTCTGCCATTCATCCGCTTATTATCACTTATTCAGGCGTAGCAACCAGGCGTTTAAGGGCACCAATAACTGCCTTAAAAAAATTACGCCCCGCCCTGCCACTCATCGCAGTACTGTTGTAATTCATTAAGCATTCTGCCGACATGGAAGCCATCACAAACGGCATGATGAACCTGAATCGCCAGCGGCATCAGCACCTTGTCGCCTTGCGTATAATATTTGCCCATAGTGAAAACGGGGGCGAAGAAGTTGTCCATATTGGCCACGTTTAAATCAAAACTGGTGAAACTCACCCAGGGATTGGCTGAGACGAAAAACATATTCTCAATAAACCCTTTAGGGAAATAGGCCAGGTTTTCACCGTAACACGCCACATCTTGCGAATATATGTGTAGAAACTGCCGGAAATCGTCGTGGTATTCACTCCAGAGCGATGAAAACGTTTCAGTTTGCTCATGGAAAACGGTGTAACAAGGGTGAACACTATCCCATATCACCAGCTCACCGTCTTTCATTGCCATACGGAACTCCGGATGAGCATTCATCAGGCGGGCAAGAATGTGAATAAAGGCCGGATAAAACTTGTGCTTATTTTTCTTTACGGTCTTTAAAAAGGCCGTAATATCCAGCTGAACGGTCTGGTTATAGGTACATTGAGCAACTGACTGAAATGCCTCAAAATGTTCTTTACGATGCCATTGGGATATATCAACGGTGGTATATCCAGTGATTTTTTTCTCCATTTTAGCTTCCTTAGCTCCTGAAAATCTCGATAACTCAAAAAATACGCCCGGTAGTGATCTTATTTCATTATGGTGAAAGTTGGAACCTCTTACGTGCCGATCACGTCTCATTTTCGCCAAAGTTGGCCAGGGCTTCCCGGTATCAACAGGGACACCAGGATTTATTTATNNTGCGAAGTGATCTTCCGTCACAGGTATTTATTCGGCGCAAAGTGCGTCGGGTGATGCTGCCAACTTACTGATTTAGTGTATGATGGTGTTTTTGAGGTGCTCCAGTGGCTTCTGTTTCTATCAGCTGTCCCTCCTGTTCAGCTACTGACGGGGTGGTGCGTAACGGCAAAAGCACCGCCGGACATCAGCGCTAGCGGAGTGTATACTGGCTTACTATGTTGGCACTGATGAGGGTGTCAGTGAAGTGCTTCATGTGGCAGGAGAAAAAAGGCTGCACCGGTGCGTCAGCAGAATATGTGATACAGGATATATTCCGCTTCCTCGCTCACTGACTCGCTACGCTCGGTCGTTCGACTGCGGCGAGCGGAAATGGCTTACGAACGGGGCGGAGATTTCCTGGAAGATGCCAGGAAGATACTTAACAGGGAAGTGAGAGGGCCGCGGCAAAGCCGTTTTTCCATAGGCTCCGCCCCCCTGACAAGCATCACGAAATCTGACGCTCAAATCAGTGGTGGCGAAACCCGACAGGACTATAAAGATACCAGGCGTTTCCCCCTGGCGGCTCCCTCGTGCGCTCTCCTGTTCCTGCCTTTCGGTTTACCGGTGTCATTCCGCTGTTATGGCCGCGTTTGTCTCATTCCACGCCTGACACTCAGTTCCGGGTAGGCAGTTCGCTCCAAGCTGGACTGTATGCACGAACCCCCCGTTCAGTCCGACCGCTGCGCCTTATCCGGTAACTATCGTCTTGAGTCCAACCCGGAAAGACATGCAAAAGCACCACTGGCAGCAGCCACTGGTAATTGATTTAGAGGAGTTAGTCTTGAAGTCATGCGCC

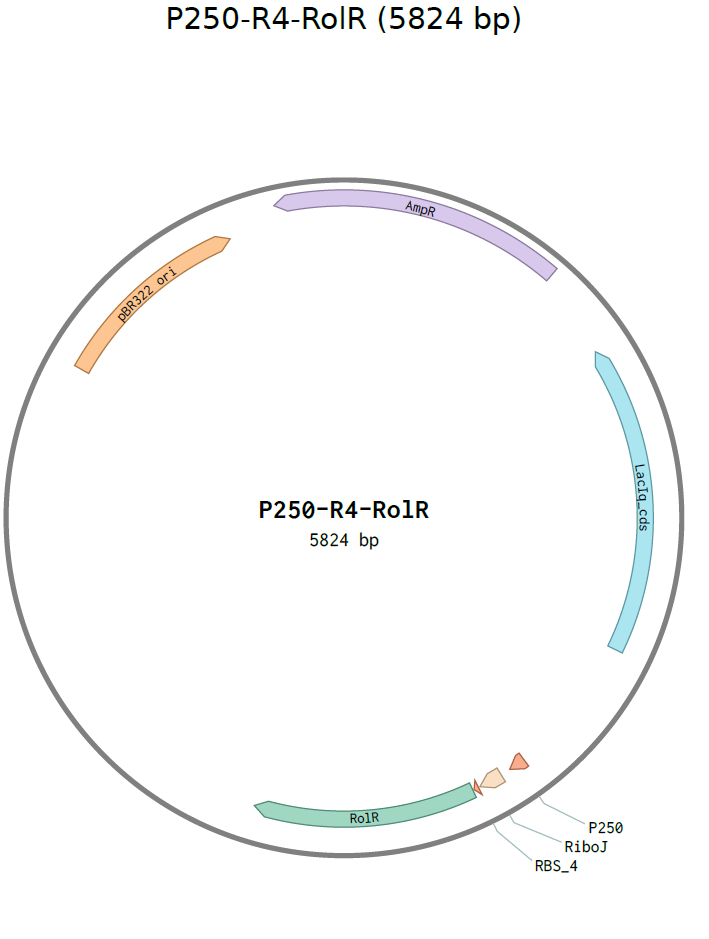

AGAAGTGGTCCTGCAACTTTATCCGCCTCCATCCAGTCTATTAATTGTTGCCGGGAAGCTAGAGTAAGTAGTTCGCCAGTTAATAGTTTGCGCAACGTTGTTGCCATTGCTGCAGGCATCGTGGTGTCACGCTCGTCGTTTGGTATGGCTTCATTCAGCTCCGGTTCCCAACGATCAAGGCGAGTTACATGATCCCCCATGTTGTGCAAAAAAGCGGTTAGCTCCTTCGGTCCTCCGATCGTTGTCAGAAGTAAGTTGGCCGCAGTGTTATCACTCATGGTTATGGCAGCACTGCATAATTCTCTTACTGTCATGCCATCCGTAAGATGCTTTTCTGTGACTGGTGAGTACTCAACCAAGTCATTCTGAGAATAGTGTATGCGGCGACCGAGTTGCTCTTGCCCGGCGTCAACACGGGATAATACCGCGCCACATAGCAGAACTTTAAAAGTGCTCATCATTGGAAAACGTTCTTCGGGGCGAAAACTCTCAAGGATCTTACCGCTGTTGAGATCCAGTTCGATGTAACCCACTCGTGCACCCAACTGATCTTCAGCATCTTTTACTTTCACCAGCGTTTCTGGGTGAGCAAAAACAGGAAGGCAAAATGCCGCAAAAAAGGGAATAAGGGCGACACGGAAATGTTGAATACTCATACTCTTCCTTTTTCAATATTATTGAAGCATTTATCAGGGTTATTGTCTCATGAGCGGATACATATTTGAATGTATTTAGAAAAATATGCGCCTTGAGCGACACGAATTATGCAGTGATTTACGACCTGCACAGCCATACCACAGCTTCCGATGGCTGCCTGACGCCAGAAGCATTGGTGCACCGTGCAGTCGATGATAAGCTGTCAAACATGAGAATTGTGCCTAATGAGTGAGCTAACTTACATTAATTGCGTTGCGCTCACTGCCCGCTTTCCAGTCGGGAAACCTGTCGTGCCAGCTGCATTAATGAATCGGCCAACGCGCGGGGAGAGGCGGTTTGCGTATTGGGCGCCAGGGTGGTTTTTCTTTTCACCAGTGAGACGGGCAACAGCTGATTGCCCTTCACCGCCTGGCCCTGAGAGAGTTGCAGCAAGCGGTCCACGCTGGTTTGCCCCAGCAGGCGAAAATCCTGTTTGATGGTGGTTAACGGCGGGATATAACATGAGCTATCTTCGGTATCGTCGTATCCCACTACCGAGATATCCGCACCAACGCGCAGCCCGGACTCGGTAATGGCGCGCATTGCGCCCAGCGCCATCTGATCGTTGGCAACCAGCATCGCAGTGGGAACGATGCCCTCATTCAGCATTTGCATGGTTTGTTGAAAACCGGACATGGCACTCCAGTCGCCTTCCCGTTCCGCTATCGGCTGAATTTGATTGCGAGTGAGATATTTATGCCAGCCAGCCAGACGCAGACGCGCCGAGACAGAACTTAATGGGCCCGCTAACAGCGCGATTTGCTGGTGACCCAATGCGACCAGATGCTCCACGCCCAGTCGCGTACCATCTTCATGGGAGAAAATAATACTGTTGATGGGTGTCTGGTCAGAGACATCAAGAAATAACGCCGGAACATTAGTGCAGGCAGCTTCCACAGCAATGGCATCCTGGTCATCCAGCGGATAGTTAATGATCAGCCCACTGACGCGTTGCGCGAGAAGATTGTGCACCGCCGCTTTACAGGCTTCGACGCCGCTTCGTTCTACCATCGACACCACCACGCTGGCACCCAGTTGATCGGCGCGAGATTTAATCGCCGCGACAATTTGCGACGGCGCGTGCAGGGCCAGACTGGAGGTGGCAACGCCAATCAGCAACGACTGTTTGCCCGCCAGTTGTTGTGCCACGCGGTTGGGAATGTAATTCAGCTCCGCCATCGCCGCTTCCACTTTTTCCCGCGTTTTCGCAGAAACGTGGCTGGCCTGGTTCACCACGCGGGAAACGGTCTGATAAGAGACACCGGCATACTCTGCGACATCGTATAACGTTACTGGTTTCACATTCACCACCCTGAATTGACTCTCTTCCGGGCGCTATCATGCCATACCGCGAAAGGTTTTGCACCATTCGATGGTGTCGGGACGTCAGGTGGCACTTTTCGGGGAAATGTGCGCGGAACCCCTATTTGTTGCGGCCGCGAAGACAGCGTTATCAGAGATGAGACACCGTGCGATAATGTCGGGCAATCAGGTGCGACTCGGTACCAAATTCCAGAAAAGAGGGGAGCGGGAAACCGCTCCCCTTTTTTCGTTTTGGTCCCAATCTATCGATTGTATGGACTTCATCTTCATTACCTCGTATCATTGTACACCTGCCGAACGAAAATATATTTTTCAAAAGTATCGTTTACCGCTAGCTCAGTCCTAGGTACAATTACAGCCATCGTACGAGCCCTGGCTGAGCACAGCTGTCACCGGATGTGCTTTCCGGTCTGATGAGTCCGTGAGGACGAAACAGCCTCTACAAATAATTTTGTTTAAACTAGCGTAACTACGGTACCACATATGCCTACCCCCTCGCAGCACAAAGACGCGAGTACTGCACAGACTGACAATCAGGTTCCTACGGGTCGTCGTGCCCAAAAGCGCGAGCAGACTCGTGCGCGTTTAATTACCAGCGCACGCACCCTGATGGCCGAACGTGGCGTGGATAATGTCGGCATTGCCGAAATCACTGAAGGTGCAAATATTGGTACTGGTACTTTTTACAATTACTTCCCCGACCGTGAACAGTTATTACAAGCTGTCGCAGAGGACGCATTCGAGTCTGTCGGTATCGCATTGGACCAGGTCTTGACGAAATTGGACGATCCTGCTGAAGTATTTGCAGGCAGCCTGCGTCACACCGTGCGCCACTCACTTGAGGATCGCATTTGGGGCGGTTTTTTCATTCAAATGGGCGCCGCGCATCCGGTGCTTATGCGTATTCTTGGACCTCGTGCACGTCGCGACTTACTTCATGGTCTGGAAACCGGCCGTTTCACTATCGAGGATTTGGACCTGGCAACAACGTGTACCTTCGGATCGCTTATTGCTGCAATTCAAATGGCGCTTAGCGCAGATCAAGATAGTAACGACGATAAAGATCAGATTTTCGCTGCTGCAATGCTTCGTATGGTTGGCGTTCAGGCCGCCGAAGCACGTGAAATCGCCTCCCGCCCGTTACCTGAAATTTCGCCCGTCAAGCCGCAGTAATAATCCTAAGAATTCGAGGAGTGCAGGCTCGGTAACATACGGTCTAGCTATCTGACTATCGCCGCTGTGAGCTCGGTACCAAATTCCAGAAAAGAGGCCGCGAAAGCGGCCTTTTTTCGTTTTGGTCCAAGCCCGATGCGCCAGAGTTGTTTCTGAAACATGGCAAAGGTATCACTAGTCTTCGCGGCCGCCATGCTGTCCAGGCAGGTAGATGACGACCATCAGGGACAGCTTCAAGGATCGCTCGCGGCTCTTACCAGCCTAACTTCGATCATTGGACCGCTGATCGTCACGGCGATTTATGCCGCCTCGGCGAGCACATGGAACGGGTTGGCATGGATTGTAGGCGCCGCCCTATACCTTGTCTGCCTCCCCGCGTTGCGTCGCGGTGCATGGAGCCGGGCCACCTCGACCTGAATGGAAGCCGGCGGCACCTCGCTAACGGATTCACCACTCCAAGAATTGGAGCCAATCAATTCTTGCGGAGAACTGTGAATGCGCAAACCAACCCTTGGCAGAACATATCCATCGCGTCCGCCATCTCCAGCAGCCGCACGCGGCGCATCTCGGGCAGCGTTGGGTCCTGGCCACGGGTGCGCATGATCGTGCTCCTGTCGTTGAGGACCCGGCTAGGCTGGCGGGGTTGCCTTACTGGTTAGCAGAATGAATCACCGATACGCGAGCGAACGTGAAGCGACTGCTGCTGCAAAACGTCTGCGACCTGAGCAACAACATGAATGGTCATCGGTTTCCGTGTTTCGTAAAGTCTGGAAACGCGGAAGTCAGCGCCCTGCACCATTATGTTCCGGATCTGCATCGCAGGATGCTGCTGGCTACCCTGTGGAACACCTACATCTGTATTAACGAAGCGCTGGCATTGACCCTGAGTGATTTTTCTCTGGTCCCGCCGCATCCATACCGCCAGTTGTTTACCCTCACAACGTTCCAGTAACCGGGCATGTTCATCATCAGTAACCCGTATCGTGAGCATCCTCTCTCGTTTCATCGGTATCATTACCCCCATGAACAGAAATCCCCCTTACACGGAGGCATCAGTGACCAAACAGGAAAAAACCGCCCTTAACATGGCCCGCTTTATCAGAAGCCAGACATTAACGCTTCTGGAGAAACTCAACGAGCTGGACGCGGATGAACAGGCAGACATCTGTGAATCGCTTCACGACCACGCTGATGAGCTTTACCGCAGCTGCCTCGCGCGTTTCGGTGATGACGGTGAAAACCTCTGACACATGCAGCTCCCGGAGACGGTCACAGCTTGTCTGTAAGCGGATGCCGGGAGCAGACAAGCCCGTCAGGGCGCGTCAGCGGGTGTTGGCGGGTGTCGGGGCGCAGCCATGACCCAGTCACGTAGCGATAGCGGAGTGTATACTGGCTTAACTATGCGGCATCAGAGCAGATTGTACTGAGAGTGCACCGGTGTGAAATACCGCACAGATGCGTAAGGAGAAAATACCGCATCAGGCGCTCTTCCGCTTCCTCGCTCACTGACTCGCTGCGCTCGGTCGTTCGGCTGCGGCGAGCGGTATCAGCTCACTCAAAGGCGGTAATACGGTTATCCACAGAATCAGGGGATAACGCAGGAAAGAACATGTGAGCAAAAGGCCAGCAAAAGGCCAGGAACCGTAAAAAGGCCGCGTTGCTGGCGTTTTTCCATAGGCTCCGCCCCCCTGACGAGCATCACAAAAATCGACGCTCAAGTCAGAGGTGGCGAAACCCGACAGGACTATAAAGATACCAGGCGTTTCCCCCTGGAAGCTCCCTCGTGCGCTCTCCTGTTCCGACCCTGCCGCTTACCGGATACCTGTCCGCCTTTCTCCCTTCGGGAAGCGTGGCGCTTTCTCATAGCTCACGCTGTAGGTATCTCAGTTCGGTGTAGGTCGTTCGCTCCAAGCTGGGCTGTGTGCACGAACCCCCCGTTCAGCCCGACCGCTGCGCCTTATCCGGTAACTATCGTCTTGAGTCCAACCCGGTAAGACACGACTTATCGCCACTGGCAGCAGCCACTGGTAACAGGATTAGCAGAGCGAGGTATGTAGGCGGTGCTACAGAGTTCTTGAAGTGGTGGCCTAACTACGGCTACACTAGAAGGACAGTATTTGGTATCTGCGCTCTGCTGAAGCCAGTTACCTTCGGAAAAAGAGTTGGTAGCTCTTGATCCGGCAAACAAACCACCGCTGGTAGCGGTGGTTTTTTTGTTTGCAAGCAGCAGATTACGCGCAGAAAAAAAGGATCTCAAGAAGATCCTTTGATCTTTTCTACGGGGTCTGACGCTCAGTGGAACGAAAACTCACGTTAAGGGATTTTGGTCATGAGATTATCAAAAAGGATCTTCACCTAGATCCTTTTAAATTAAAAATGAAGTTTTAAATCAATCTAAAGTATATATGAGTAAACTTGGTCTGACAGTTACCAATGCTTAATCAGTGAGGCACCTATCTCAGCGATCTGTCTATTTCGTTCATCCATAGTTGCCTGACTCCCCGTCGTGTAGATAACTACGATACGGGAGGGCTTACCATCTGGCCCCAGTGCTGCAATGATACCGCGCGATCCACGCTCACCGGCTCCAGATTTATCAGCAATAAACCAGCCAGCCGGAAGGGCCGAGCGC
